# Fundamental limits of amplification-free CRISPR-Cas12 and Cas13 diagnostics

**DOI:** 10.1101/2022.01.31.478567

**Authors:** Diego A. Huyke, Ashwin Ramachandran, Vladimir I. Bashkirov, Efthalia K. Kotseroglou, Theofilos Kotseroglou, Juan G. Santiago

## Abstract

Interest in CRISPR diagnostics continues to increase. CRISPR-Cas12 and -Cas13 based detection are particularly interesting as they enable highly specific detection of nucleic acids. The fundamental sensitivity limits of Cas12 and Cas13 enzymes are governed by their kinetic rates and are critical to develop amplification-free assays. However, these kinetic rates remain poorly understood and their reporting has been inconsistent. We here measure kinetic parameters for several enzymes (LbCas12a, AsCas12a, AapCas12b, LwaCas13a and LbuCas13a) and evaluate their limits of detection (LoD) for amplification-free target detection. Collectively, we here present quantitation of enzyme kinetics for 14 gRNAs and nucleic acid targets for a total of 50 sets of kinetic rate parameters and 25 LoDs. Importantly, we also validate the self-consistency our measurements by comparing trends and limiting behaviors with a Michaelis-Menten, trans-cleavage reaction kinetics model. Our measurements reveal that activated Cas12 and Cas13 enzymes exhibit typical trans-cleavage catalytic efficiencies between order 10^5^ and 10^6^ M^-1^ s^-1^. Moreover, for assays that use fluorescent reporter molecules (ssDNA and ssRNA) for target detection, we find most CRISPR enzymes have an amplification-free LoD in the picomolar range. We find also that successful detection of target requires cleavage (by activated CRISPR enzyme) of at least ~0.1% of the fluorescent reporter molecules. This fraction of cleaved reporters is required to differentiate signal from background, and we hypothesize that this fraction is largely independent of the detection method (i.e., endpoint vs reaction velocity). Our results provide a map of the feasible application range and highlight areas of improvement for the emerging field of CRISPR diagnostics.

## 1. INTRODUCTION

CRISPR diagnostics have received intense attention over the last several years.^1^ Amplification-free CRISPR diagnostics, specifically, seek to offer specific and sensitive nucleic acid detection at the point of care.^2–6^ These assays leverage programmable subtypes of Cas12 and Cas13 to respectively identify DNA and RNA directly in samples. However, the catalytic rates of these enzymes and, therefore, the achievable LoDs of these assays remains an open question. We recently demonstrated that many published reports quantifying kinetic rates of Cas12 and Cas13 exhibit serious violations of species conservation principles and inconsistent reaction rate limits.^7–11^

We chose to study Cas12 and Cas13 here as they are ubiquitous in CRISPR diagnostics. LbCas12a is the most widely used enzyme to detect DNA targets.^8,12–14^ Its reported turnover rates (*k_cat_*) have varied from 0.069 to 4850 s^-1^ while Michaelis-Menten constants (*K_M_*) have varied from 13 to 1000 nM.^5,7,9,15^ Hence, interestingly, reported catalytic efficiencies (*k_cat_/K_M_*) for this enzyme have varied by more than four orders of magnitude. For AsCas12a, Cofsky *et al*. is the only study which has reported enzyme kinetic parameters (*k_cat_* = 0.6 s^-1^ and *K_M_* = 2700 nM).^16^ AapCas12b is an attractive Cas12 subtype because it exhibits trans-cleavage at temperatures compatible with LAMP (loop mediated isothermal amplification).^17^ However, no kinetic parameters have been reported for this enzyme. For Cas13-based detection, LbuCas13a has been widely used and reported *k_cat_* values have varied from 3.3 to 4900 s^-1^, while *K_M_* values have varied from 760 to 4500 nM.^5,18^ LwaCas13a is another commonly used Cas13 enzyme, but we know of no reported characterization of its kinetics.

In brief, there has been no self-consistent study of the kinetic rate parameters of Cas12 and Cas13 subtypes combined with an effort to relate these kinetic rates to achievable LoDs. To guide future work, we rigorously characterize fundamental detection limits and estimate the currently achievable sensitivity of CRISPR diagnostics. We first report Michaelis-Menten parameters for LbCas12a, AsCas12a, and AapCas12b using 10 different gRNAs and 20 different targets. We then supplement these data with measured LoDs for each Cas-gRNA-target combination. We further apply the same procedure to LbuCas13a and LwaCas13a using four different gRNA-target pairs. We use an experimentally validated enzyme kinetics model^7^ to demonstrate the self-consistency of our experimental data. Lastly, we use the measured rate parameters and the model to map an operational space for CRISPR diagnostics in terms of enzyme trans-cleavage catalytic efficiency, LoD, fraction of reporters cleaved, and assay time. Our findings may also help guide the design of CRISPR assays that rely on nucleic acid amplification by, for example, providing an estimate of amount of amplification required prior to CRISPR-based detection.

## 2. RESULTS

### 2.1 Cas12 Michaelis-Menten kinetics

We first measured kinetic parameters for LbCas12a, AsCas12a, and AapCas12b. We complexed these enzymes with gRNAs complementary to ss- and dsDNA targets (**Fig. 1a**). Complexed enzymes that recognized targets then collaterally cleaved fluorophore-quencher ssDNA reporters. This yielded a fluorescence increase at a rate governed by *k_cat_* and *K_M_* (see **Methods** for complete experimental protocol). Importantly, *k_cat_* and *K_M_* depend on the gRNA, target, and enzyme subtype and are thus not fixed for a given Cas enzyme.^7^

**Fig. 1.**
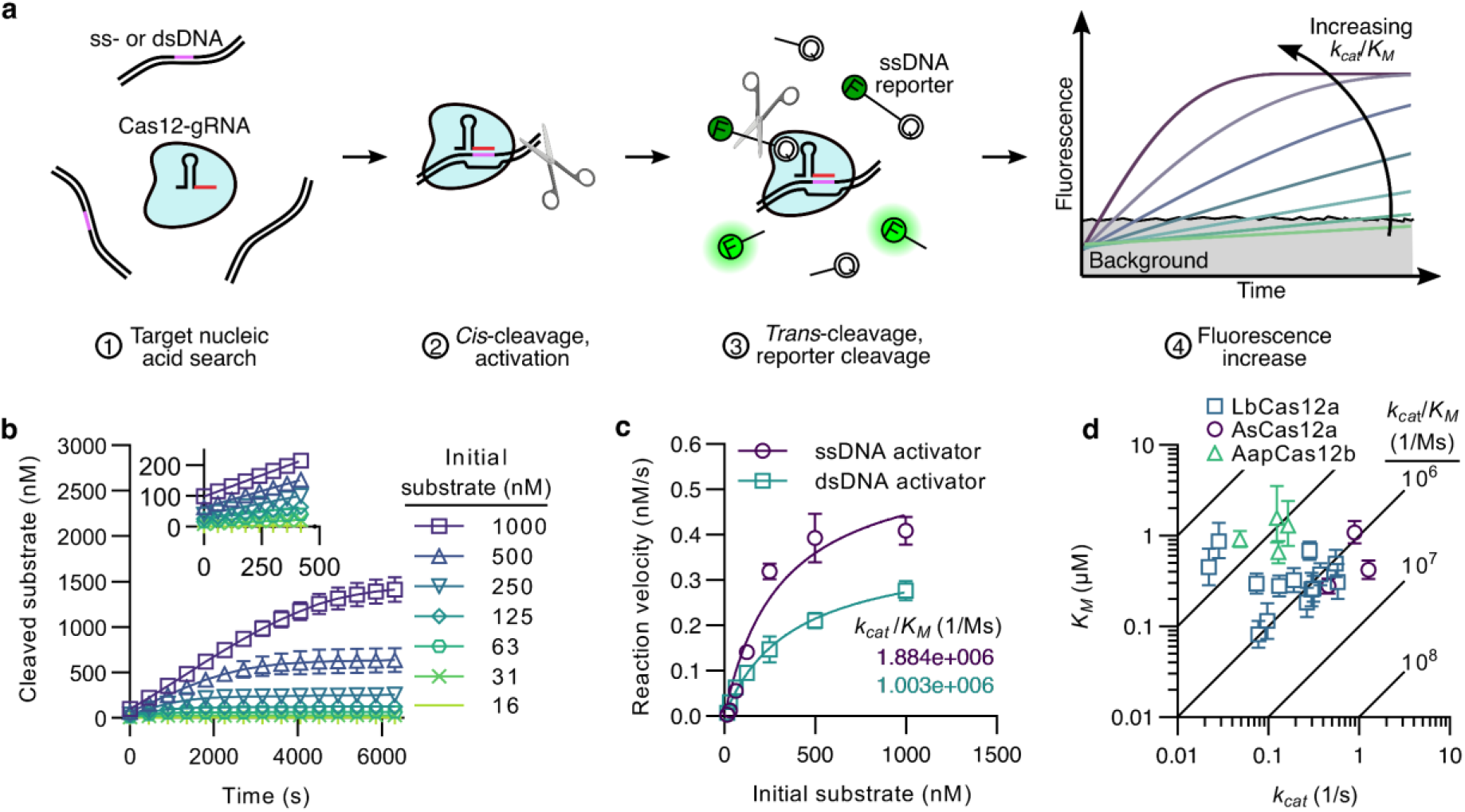
Michaelis-Menten kinetics of CRISPR-Cas12. **a** Schematic for ss- or dsDNA detection using a Cas12-gRNA complex. Upon recognition of the target DNA, the enzyme initiates collateral cleavage of ssDNA reporters, increasing fluorescence signal. For a fixed target concentration and typical assay conditions,^7^ the rate of increase in fluorescence is governed by the catalytic efficiency *k_cat_/K_M_* of the Cas12-gRNA-activator complex. **b** An example set of progress curves of cleaved substrate concentration versus time. Data shown here is for LbCas12a complexed with gRNA 1 (termed LbCas12a-1) and activated with 1 nM of dsDNA target. The inset shows the linear portion of the curve. **c** Initial reaction velocity versus substrate concentration for LbCas12a-1 activated with ss- or dsDNA (corresponding to the data in **b**). A nonlinear fit to a Michaelis-Menten curve was performed to obtain (for ss- and dsDNA, respectively) *k_cat_* = 0.58 and 0.38 s^-1^ and *K_M_* = 310 and 370 nM. **d** 21 different measurements of *K_M_* versus *k_cat_* across LbCas12a, AsCas12a, and AapCas12b enzymes complexed with 10 gRNAs and 20 activators. The diagonal lines represent axes of constant *k_cat_/K_M_*. Note most enzymes have *k_cat_/K_M_* on the order of 10^5^ to 10^6^ M^-1^ s^-1^.

We varied LbCas12a collateral cleavage at reporter concentrations between 16 and 1000 nM for a 1 nM amount of target activator (**Figs. 1b** and **S1**). The data shown here is for LbCas12a complexed with gRNA 1 (hereafter termed LbCas12a-1) and activated with the corresponding dsDNA target (sequences in **Tables S2-S4**). The set of initial reaction velocities for each reactant system was used to fit a Michaelis-Menten curve and estimate *k_cat_* and *K_M_* for LbCas12a-1 activated by ss- or dsDNA targets (**Fig. 1c**).

We obtained Michaelis-Menten curves for several other LbCas12a, AsCas12a, and AapCas12b trans-cleavage reactions (**Figs. S2-S5**). Interestingly, *k_cat_* and *K_M_* exhibited no discernable trends for the same Cas12-gRNA complex activated by ss-versus dsDNA (**Table 1** and **S1**). That is, unlike previous reports, dsDNA targets did not necessarily yield higher *k_cat_/K_M_* than ssDNA targets.^8^ Contrary to the case of a dsDNA target, we found that a ssDNA target did not measurably activate AsCas12a-1 (**Fig. S4**), which is also in contrast with previous observations.^19^ Furthermore, AapCas12b showed no significant change in *k_cat_/K_M_* at 37 versus 60 °C (**Fig. S5**). This suggests AapCas12b can effectively detect targets over a temperature range inclusive of most isothermal nucleic acid amplification reactions (e.g., LAMP).^17^

**Table 1.**
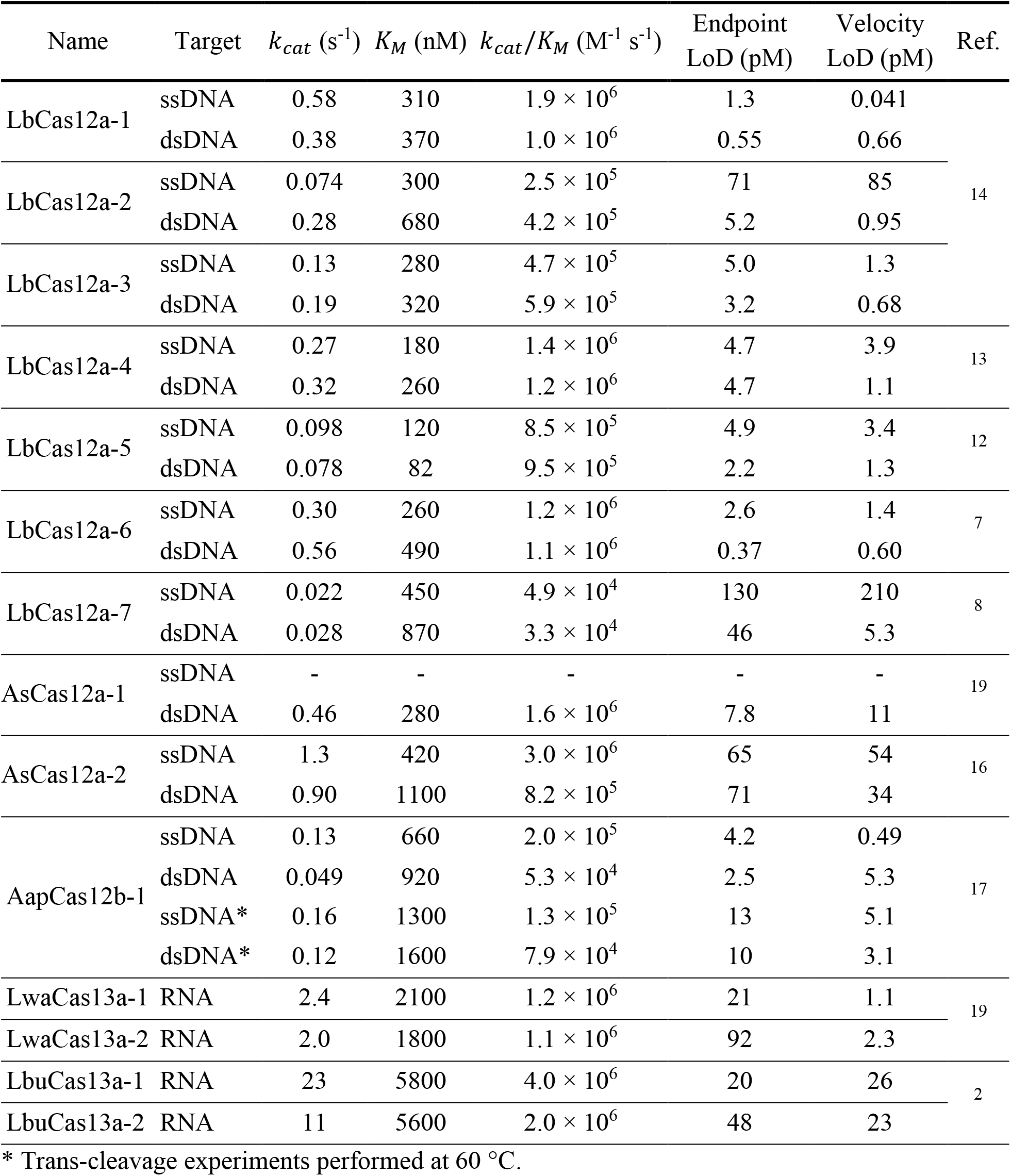
Summary of experimentally measured kinetic parameters and corresponding LoDs based on both endpoint and maximum velocity detection methods.

Taken together, our experiments with these three Cas12 enzymes, including a total of 21 different trans-cleavage reactions (**Fig. 1d**), revealed a range of *k_cat_* values between order 0.01 and 1 s^-1^ and *K_M_* values between order 100 and 1000 nM. Importantly, *k_cat_/K_M_* values were on the order of 10^5^ to 10^6^ M^-1^ s^-1^. We found that our measured *k_cat_/K_M_* are 1 to 3 orders of magnitude lower than most reported values^5,9,13,15^ (see also a summary^7^ of trends in reported values).

### 2.2 Cas12 limits of detection and relation to kinetics

Next, we measured LoDs for LbCas12a, AsCas12a, and AapCas12b. Herein, we first observed LbCas12a-1 collateral cleavage over 1 h in the presence of a dsDNA target (between 10^-2^ and 10^4^ pM) as well as in the absence of a target (no template control, NTC, **Fig. 2a**). We defined and quantified an LoD concentration based on endpoint fluorescence (*L_E_*) or maximum reaction velocity (*L_V_*). Note *L_E_* and *L_V_* have units of concentration and are derived from raw fluorescence signals (see **Methods**). In both schemes, any measurement statistically distinguishable (by 3 standard deviations) from the NTC corresponded to a detected target (see dashed lines in **Figs. 2b** and **2c**). Importantly, *L_V_* follows from measurements of non-specific Cas-gRNA trans-cleavage activity (hereafter termed background activity) and/or reporter degradation, each of which has been reported for CRISPR systems.^2,4,5,13,15,18,20^

**Fig. 2.**
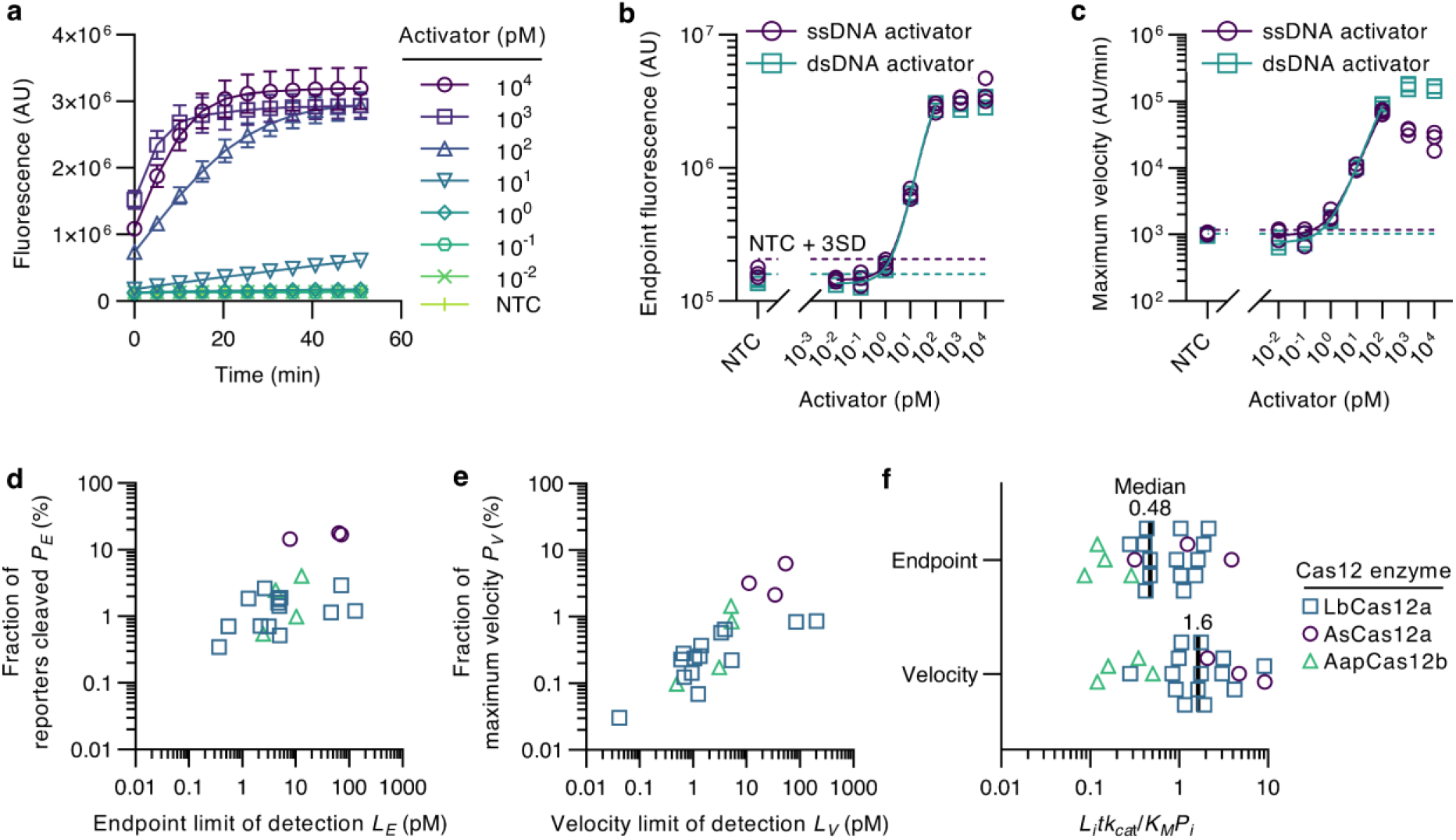
LoD studies for CRISPR-Cas12. **a** Fluorescence versus time of LbCas12a-1 for fixed (initially uncleaved) reporter concentration of 200 nM and dsDNA target concentrations between 10^-2^ and 10^4^ pM. **b** Endpoint fluorescence for LbCas12a-1 activated by ss- or dsDNA target. The dashed lines correspond to the detection threshold. The solid line is a four-parameter logistic (4PL) curve fit to the experimental data which was used to estimate an endpoint LoD of 1.3 (for ssDNA) and 0.55 (for dsDNA) pM. **c** Maximum velocity of progress curves for LbCas12a-1 activated by varied ss- or dsDNA target concentrations. The estimated velocity LoD is 0.041 (for ssDNA) and 0.66 (for dsDNA) pM. **d** Fraction of cleaved reporters at the endpoint LoD (*P_E_*) versus endpoint LoD (*L_E_*) for various LbCas12a, AsCas12a, and AapCas12b gRNA-target combinations. **e** Fraction of maximum cleavage velocity across all targets and concentrations (*P_V_*) versus velocity LoD (*L_V_*) for the same Cas12-gRNA-target combinations. **f** Heuristic *H* = *L_i_tk_cat_/K_M_P_i_* evaluated for *i* = *E* (endpoint) and *V* (velocity). Our analysis suggests this heuristic should be order unity when the measured LoD approaches the predicted LoD for a given Cas12-gRNA-target catalytic efficiency.

The latter experiments were repeated for all Cas12-gRNAs studied here (**Figs. S6-S9**). The observed endpoint LoDs varied between 0.37 and 130 pM. Meanwhile, velocity LoDs varied more significantly between 0.041 and 210 pM, likely reflecting the variability in the non-specific activity among Cas-gRNA-target combinations. AsCas12a exhibited the highest background activity (**Fig. S8**). For example, the median AsCas12a background activity was 3-fold higher than the median LbCas12a background activity across the reactions studied here. We also found that velocity-based detection improved the LoD (relative to endpoint detection) by up to an order of magnitude for cases of low background activity (**Table 1**). The results strongly highlight the importance of background enzyme activity on CRISPR assay sensitivity.

We next explored the relation between *L_E_* and the fraction of reporters cleaved (*P_E_*) when the target concentration equaled *L_E_* (**Fig. 2d**). This explores a hypothesis that *L_E_* scales with *P_E_*, and that *P_E_* is limited by the enzyme background activity and signal from free dye. Analogously, we compared *L_V_* and the ratio of the maximum reaction velocity when the target concentration equaled *L_V_* and the maximum reaction velocity across target concentrations (*P_V_***, Fig. 2e**). The background associated with *L_E_* is likely influenced by imperfectly quenched reporters, degraded reporters, background enzyme activity, and/or free dye, and this may explain the large degree of scatter in this *P_E_* plot. Overall, we found that *P_E_* was typically higher than *P_V_*, which suggests that velocity-based detection is less sensitive to irreproducibility in the background signal of reporter solutions.

Lastly, we explored the overall agreement between measured kinetic parameters and LoDs (**Fig. 2f**) using a heuristic quantity *H* defined as

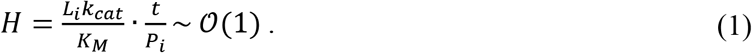

Here, *t* is the duration of the LoD experiment (set here to 1 h), *L_i_* is either *L_E_* or *L_V_*, *k_cat_/K_M_* is the catalytic efficiency of the activated Cas12-gRNA complex, and *P_i_* is either *P_E_* or *P_V_*. Conceptually, *K_M_/L_E_k_cat_* is the reaction time scale when the substrate concentration is less than *K_M_* and therefore *L_E_k_cat_/K_M_t* is a heuristic for the degree of completion of the trans-cleavage reaction.^7^ We hypothesize that this degree of completion scales with the measured fraction of cleaved reporters *P_E_*. Though the heuristic *H* is strictly applicable only for endpoint fluorescence measurements (*L_E_* and *P_E_*), we here posit a similar scaling for velocity-based measurements (comparing *L_V_* and *P_V_*) as well. For self-consistency, *H* should be near order unity. Importantly, our heuristic relates the enzyme kinetic properties with observed LoDs across Cas12 subtypes, gRNAs, and targets. Across our Cas12 data, we found all values of *H* were within an order of magnitude of unity, suggesting consistency between measured kinetics and observed LoDs.

### 2.3 Cas13 Michaelis-Menten kinetics and limit of detection

Following, we measured kinetic parameters and LoDs for LwaCas13a and LbuCas13a. We measured LwaCas13a-1 collateral cleavage at reporter concentrations between 125 and 6000 nM for a 1 nM amount of target activator (**Fig. 3a**). At these reporter concentrations, the inner filter effect^21^ required a non-linear standard curve (**Fig. S10**). Michaelis-Menten curves were then fit to data for LwaCas13a-1 and −2 (**Figs. 3b** and **S11**) in addition to LbuCas13a (**Fig. S12**). Our Cas13 experiments (**Fig. 3c**) revealed *k_cat_* values between order 1 and 10 s^-1^ and *K_M_* values of order 1000 nM. The absolute values of *k_cat_* and *K_M_* were noticeably higher for the Cas13 systems than for the Cas12 systems, but *k_cat_/K_M_* nevertheless remained on the order of 10^6^ M^-1^ s^-1^. Again, our measured *k_cat_/K_M_* is at least 1 to 3 orders of magnitude lower than reported values.^2,5,18,22^

**Fig. 3.**
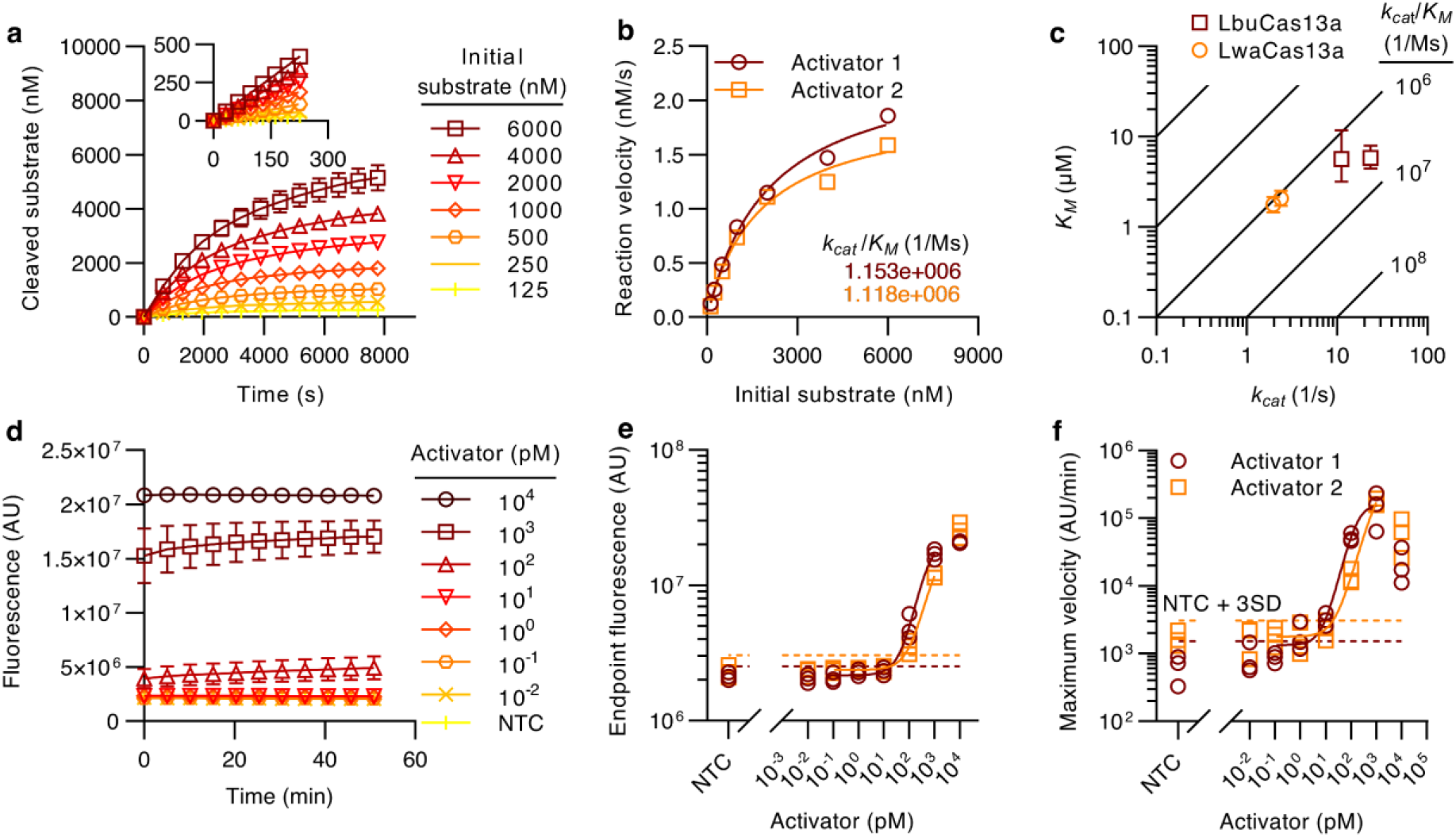
Michaelis-Menten and LoD studies for CRISPR-Cas13. **a** Progress curves of cleaved substrate versus time for LwaCas13a-1. Data shown here is for a fixed activated enzyme concentration of 1 nM. The inset shows the linear portion of the curve. **b** Initial reaction velocity versus substrate concentration for LwaCas13a-1 and −2. A nonlinear fit to a Michaelis-Menten curve was performed to obtain (for LwaCas13a-1 and −2, respectively) *k_cat_* = 2.4 and 2.0 s^-1^ and *K_M_* = 2100 and 1800 nM. **c** *K_M_* versus *k_cat_* for LbuCas13a and LwaCas13a, each complexed with two gRNAs. The diagonal lines represent axes of constant *k_cat_/K_M_*. Although these Cas13 enzymes have higher *k_cat_* and *K_M_* relative to Cas12, their *k_cat_/K_M_* were also below about 10^7^ M^-1^ s^-1^. **d** Fluorescence signal versus time of LwaCas13a-1 for fixed (initially uncleaved) reporter concentration of 1 μM and RNA target concentrations between 10^-2^ and 10^4^ pM. **e** Endpoint fluorescence for LwaCas13a-1 and −2 as a function of RNA target concentration. The dashed lines correspond to the detection threshold. The solid line is a 4PL curve fit to the experimental data which was used to estimate an endpoint LoD of 21 and 92 pM (for LwaCas13a-1 and −2, respectively). **f** Maximum velocity of progress curves for LwaCas13a-1 and −2 as a function of RNA target concentration. The estimated velocity LoD is 1.1 and 2.3 pM (for LwaCas13a-1 and −2, respectively).

We lastly measured LoDs for LwaCas13a and LbuCas13a. We showed collateral cleavage by LwaCas13a-1 over 1 h when activated by an RNA target between 10^-2^ and 10^4^ pM (**Fig. 3d**). Endpoint- and velocity-based LoDs were calculated for LwaCas13a-1 and −2 (**Figs. 3e-f** and **S13**) as well as LbuCas13a (**Fig. S14**). Velocity-based detection improved the LoD by one order of magnitude in half of the cases (**Table 1** and **Fig. S14d**). Most importantly, the Cas13 and Cas12 LoDs were on the same order. We finalized our Cas13 analysis by applying our heuristic method (**Fig. S15**), and this showed a moderate correlation between *k_cat_/K_M_* and LoDs.

## 3. DISCUSSION

After exploring kinetic parameters and LoDs for Cas12 and Cas13, we finally predicted a feasible sensitivity range for CRISPR diagnostics. Recall *K_M_/Lk_cat_* (where *L* is the target concentration) is the time required for activated enzymes to cleave ~63% of reporters.^7^ However, as we showed, high-quality detectors and Cas-gRNA complexes can positively identify the presence of a target when as few as 0.3% of reporters are cleaved (**Fig. 2d**). We summarize these observations and analysis by plotting an approximate relationship between achievable LoD and the required catalytic efficiency (**Fig. 4**). We show the dependence of this relation over a wide variety of assay times and fractions of cleaved reporters *P*. We vary *P* by 7 orders of magnitude but stress that our measurements suggest *P* should be at least above 0.1% given the fold-change in the quenched versus unquenched fluorescence signal of current reporter molecules.

**Fig. 4.**
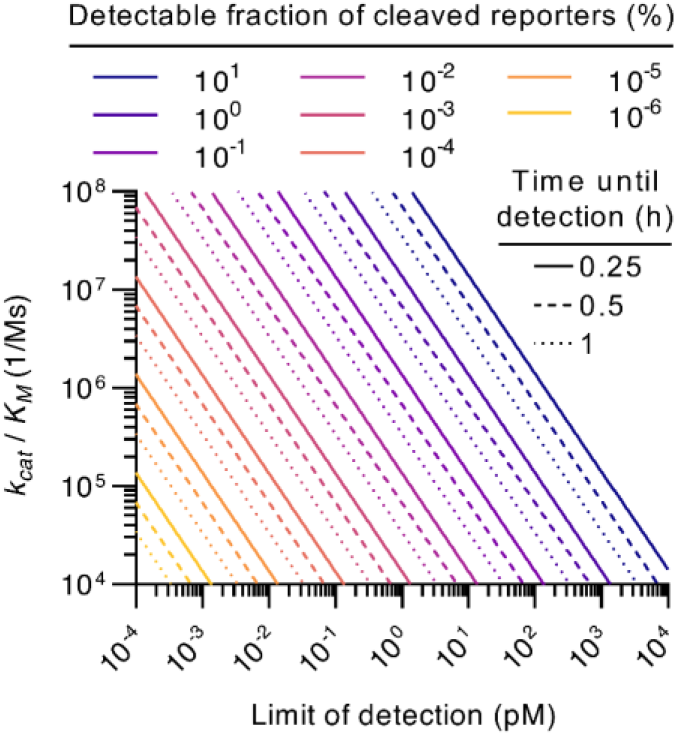
Catalytic efficiency *k_cat_/k_M_* versus predicted LoD for cleaved reporter fractions between 10^-5^ and 10%. Data is shown for assay times between 0.25 and 1 h. The plot was generated by fixing *K_M_* = 200 μM and the initial reporter concentration (= 200 nM). Note that, in this work, we found *k_cat_/k_M_* mostly between 10^5^ and 10^6^ M^-1^ s^-1^. Given a detectable fraction of cleaved reporters of 0.1%, the applicability of CRISPR diagnostics is therefore limited to target concentrations above or near 1 pM.

Our findings indicated the LoD of CRISPR diagnostics is largely governed by *k_cat_/K_M_*, *P*, and the detection time. For example, given *P* = 1% and a time of 1 h, a 10-fold improvement in *k_cat_/K_M_* (*e.g*., 10^5^ to 10^6^ M^-1^ s^-1^) provides a 10-fold improvement in LoD (30 to 3 pM). Similarly, given *k_cat_/K_M_* = 10^6^ M^-1^ s^-1^ and a time of 1 h, a 10-fold improvement in *P* (*e.g*., from 1 to 0.1%) improves the LoD 10-fold (3 to 0.3 pM). Moreover, detection time is also proportional to LoD (*e.g*., 60 versus 15 min improves LoD by 4-fold). Experiments much longer than 1 h, however, compromise the desirability of CRISPR over other techniques. LoDs could be improved with increased reporter dynamic range (i.e., cleaved-to-uncleaved fluorescence intensity ratio). We here found ratios of respectively 9.7 (**Fig. S1**) and 4.9 (**Fig. S10**) for our ssDNA and ssRNA reporters. Another way to increase LoD is to preconcentrate the target concentration and thereby increase the reaction rate. For example, new droplet based techniques may enable target detection in the single femtomolar range (or below) by distributing target molecules within femto- or picoliter droplets, thereby locally increasing target concentration within the droplets.^4,15^ As another example, electrokinetic preconcentration methods can also be used to increase the concentration of target by ~1,000-fold within a reaction volume.^12^

Interestingly, the LoDs reported by, for example, Fozouni *et al*.^2^ are surprising given our data and analysis. Given the time, catalytic efficiency, and LoD reported therein, the system of Fozouni *et al*. reportedly detects a signal distinguishable from background when only order 0.001% of reporters have been cleaved. Finally, we note that our model and assay design principles may be broadly applicable to other detection methods such as those which rely on electrochemical impedance measurements for target detection.^23–25^

## METHODS

### Enzyme preparation

LbCas12a was purchased from New England Biolabs (NEB, MA, USA) at a concentration of 100 μM. AsCas12a and LwaCas13a were obtained from IDT at concentrations, respectively, of 64 and 72 μM. The plasmids for AapCas12b (pAG001 His6-TwinStrep-SUMO-AapCas12b) were a gift from Omar Abudayyeh and Jonathan Gootenberg (http://n2t.net/addgene:153162; RRID: Addgene_153162). The plasmids for LbuCas13a (pC0072 LbuCas13a His6-TwinStrep-SUMO-BsaI) were a gift from Feng Zhang (http://n2t.net/addgene:115267; RRID: Addgene_115267). Expression plasmids were transformed into Rosetta 2(DE3). pLysS competent cells (Millipore Corporation, MA, USA) and protein expression was performed in 2 L of Terrific Broth supplemented with 50 μg mL^-1^ carbenicillin (Teknova Inc., CA, USA) at 18 °C for 16 hours. Proteins were purified according to the published protocol^26^ without major modification except the harvested cells were lysed by sonication. Proteins were stored in Storage Buffer (600 mM NaCl, 50 mM Tris-HCl pH 7.5, 5% glycerol, 2mM DTT) at −80 °C in 10 μl aliquots. After purification, the stock AapCas12b and LbuCas13a enzyme concentrations were respectively 4.4 and 39 μM.

### Trans-cleavage kinetics experiments

To measure the trans-cleavage kinetics of Cas12 enzymes, we first prepared 1 μM solutions of the Cas12-gRNA complex. This was achieved by incubating a mixture of 1 μM synthetic gRNA with at least 2-fold excess of the corresponding Cas12 at 37 °C for 30 min on a hot plate. Cas12-gRNA complexes were then activated for trans-cleavage activity by mixing and incubating these complexes with synthetic ss- or dsDNA at 100 nM concentration at 37 °C for 30 min on a hot plate. The latter step yielded a solution with an activated Cas12 concentration of 100 nM. We performed the trans-cleavage kinetics assay using 1 nM of activated Cas12 and varied ssDNA reporter concentrations of 15.63, 31.25, 62.5, 125, 250, 500, 1000, and 2000 nM. Three replicates were taken for each concentration. All LbCas12a and AsCas12a reactions were buffered in 1x NEBuffer 2.1 (composed of 50 mM NaCl, 10 mM Tris-HCl, 10 mM MgCl2, and 100 μg mL^-1^ bovine serum albumin at pH = 7.9) and all AapCas12b reactions were buffered in 1x isothermal amplification buffer (composed of 20 mM Tris-HCl, 10 mM (NH_4_)_2_SO_4_, 50 mM KCl, 2 mM MgSO_4_, 0.1% Tween® 20 at pH 8.8) supplemented with 10 mM MgCl_2_.^17^ Most Cas12 trans-cleavage reactions were run at 37 °C (AapCas12b was reacted at 37 and 60 °C). Cas12 kinetics data was measured using a MiniOpticon thermal cycler (Bio-Rad Laboratories, CA, USA), and fluorescence was measured every 30 s. A fluorescence calibration curve (**Fig. S1**) was used to convert fluorescence values (in arbitrary units) to molar concentrations (in nM). The initial reaction velocities (in nM s^-1^) for Michaelis-Menten analyses were obtained using a linear fit to the first ~300 s of the reaction progress curves. The reaction velocities versus reporter concentration data were fitted to the Michaelis-Menten equation using GraphPad Prism 9 (GraphPad Software, CA, USA) to obtain *k_cat_* and *K_M_*.

The Cas13 trans-cleavage assay was like that followed for Cas12 with the following differences. First, Cas13-gRNA was activated using synthetic ssRNA targets. The reporter ssRNA concentrations used were 125, 250, 500, 1000, 2000, 4000, and 6000 nM. The buffer used for all Cas13 reactions consisted of 20 mM HEPES, 50 mM NaCl, 10 mM MgCl2, RNase inhibitor 1 U mL^-1^, and 5% glycerol at pH ≈ 6.8. Cas13 kinetics data was measured using an Infinite 200 PRO plate reader (Männedorf, Switzerland), and fluorescence was measured every 30 s. Due to the necessary high reporter concentrations, an inner filter effect was observed which reduced the apparent fluorescence of reporters. A power law relation was therefore used to convert fluorescence values to molar concentrations (Fig. S10).^21^ The initial reaction velocities (in nM s^-1^) for Michaelis-Menten analyses were obtained using a linear fit to the first ~200 s of the reaction progress curves.

### Limit of detection experiments

To determine the LoD for each gRNA-target pair for both Cas12 and Cas13 enzymes, we first prepared a solution containing 1 μM Cas-gRNA complex using the procedure described for the trans-cleavage assay. Later, individual reactions were set up by mixing Cas-gRNA complex, reporter ssDNA or ssRNA, and varying amounts of target activator (ssDNA/dsDNA/RNA), for each enzyme in the corresponding reaction buffer. For this, we used 100 nM of Cas-gRNA, 200 nM and 1 μM reporters for Cas12 and Cas13 experiments respectively and varied the target activator concentration between 10 nM and 10 fM in 10-fold dilutions. Fluorescence readouts were taken in 1 min intervals for a total of 60 min on the ABI 7500 Fast thermal cycler (Applied Biosystems, CA, USA). LoDs were estimated using two methods: endpoint fluorescence readout and maximum reaction velocity (i.e., slope). For the former method, we used the fluorescence readout at 60 min as the measure of signal. For the maximum reaction velocity method, we used the maximum value of instantaneous reaction velocity over the course of the experiment (obtained from the slope of fluorescence versus time, averaged over 3 min) as a measure of signal. A 4PL curve was then fit to the data for each enzyme and method.

### Oligo preparation

ssDNA and RNA oligonucleotides (target activator, gRNA, and reporter) were purchased from Integrated DNA Technologies (IDT, IA, USA), Elim Biopharmaceuticals Inc. (CA, USA), and GeneLink Inc. (FL, USA). RNA oligonucleotides (from IDT and GeneLink) were resuspended to 100 μM in RNA reconstitution buffer (GeneLink). DNA oligonucleotides (Elim) were obtained at 100 μM concentration in nuclease-free water. The complete list of oligos used in this study is provided (**Tables S2-S4**). For Cas12 experiments, synthetic dsDNA target oligos were prepared by hybridizing 10 μM of ssDNA template with 50 μM of complementary ssDNA in 1x NEBuffer 2.1 (NEB), to provide a solution with an effective dsDNA concentration of 10 μM. The hybridization protocol included a 95 °C hold for 2 min, followed by a slow cool (0.1 °C s^-1^) to 25 °C. The protocol was carried out using the MiniOpticon thermal cycler (Bio-Rad).

### Calculation of LoD

This section describes the calculation of the endpoint-based (*L_E_*) and velocity-based (*L_V_*) LoDs. Specifically, *L_E_* was quantified as the molar concentration of the target at which the measured endpoint fluorescence signal equaled the endpoint threshold. The endpoint threshold *T_E_* was defined as

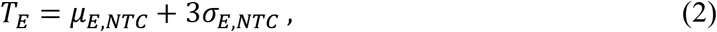

where *μ_E,NTC_* and *σ_E,NTC_* respectively correspond to the mean and standard deviation of the endpoint fluorescence measurements of the NTC samples. Analogously, *L_V_* was based on the maximum observed reaction velocity and its comparison to the background, non-specific activity of the enzyme complex with fluorescence probes and/or probe degradation (i.e., the NTC cleavage activity). The analogous, velocity-based threshold *T_V_* was defined as

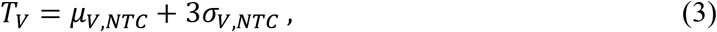

where *μ_V,NTC_* and *σ_V,NTC_* respectively correspond to the mean and standard deviation of the maximum cleavage (measured) velocities of the NTC samples.

## ACKNOWLEDGEMENTS

D.A.H. is supported by a National Science Foundation Graduate Research Fellowship. A.R. acknowledges support from the Bio-X Bowes Fellowship of Stanford University. All authors gratefully acknowledge support Ford Motor Company and the Stanford Bio-X Interdisciplinary Initiatives Committee (IIP).

## AUTHOR CONTRIBUTIONS

D.A. Huyke and A. Ramachandran, and J.G. Santiago conceived of the study. D.A. Huyke and A. Ramachandran performed the Michaelis-Menten and LoD experiments. D.A. Huyke, A. Ramachandran, and J.G. Santiago wrote the manuscript. V.I. Bashkirov, E.K. Kotseroglou, and T. Kotseroglou prepared the enzymes and contributed to data analysis. All authors have discussed the results and the interpretation.

## COMPETING INTERESTS

The authors declare no competing interests.

## Notes

### Competing Interest Statement

The authors have declared no competing interest.

