## Supplementary information for "Fundamental limits of amplification-free CRISPR-Cas12 and Cas13 diagnostics"

**Contents**

S1. DNA reporter calibration curves
S2. LbCas12a, AsCas12a, and AapCas12b Michaelis-Menten kinetics
S3. LbCas12a, AsCas12a, and AapCas12b limits of detection
S4. RNA reporter calibration curves
S5. LwaCas13a and LbuCas13a Michaelis-Menten kinetics
S6. LwaCas13a and LbuCas13a limits of detection
S7. Cas13a correlations
S8. Michaelis-Menten kinetic parameters with confidence intervals
S9. Lists of oligos used in this work

### S1. DNA reporter calibration curves

**Figure S1** shows the linear calibration curves for the cleaved and uncleaved ssDNA reporters used in this work. The calibration curves were used to quantify cleavage activity in molar units.[1]

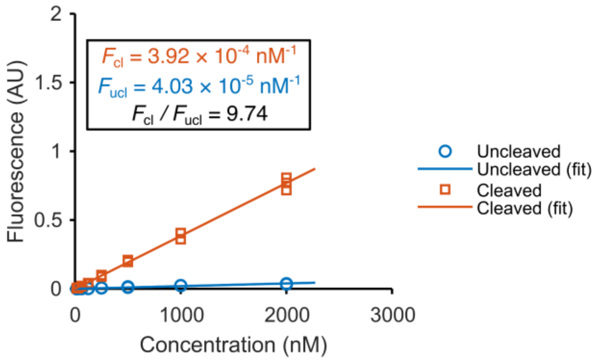

**Fig. S1. DNA reporter calibration curves.** Fluorescence versus concentration of uncleaved and cleaved FAM-BHQ DNA reporters. Lines of best fit to the experimental data (solid lines) were obtained by linear regression. $F_{ucl}$ and $F_{cl}$ are the slopes of the fits to, respectively, the uncleaved and cleaved fluorescence data. Data was taken in a MiniOpticon™ System (Bio-Rad Laboratories, Inc., USA).

### S2. LbCas12a, AsCas12a, and AapCas12b Michaelis-Menten kinetics

This section presents the linear portion of the measured trans-cleavage progress curves for the LbCas12a, AsCas12a, and AapCas12b complexes (**Figs. S2-S5**). Also presented are the Michaelis-Menten fits to the initial reaction velocity to extract relevant kinetic parameters.

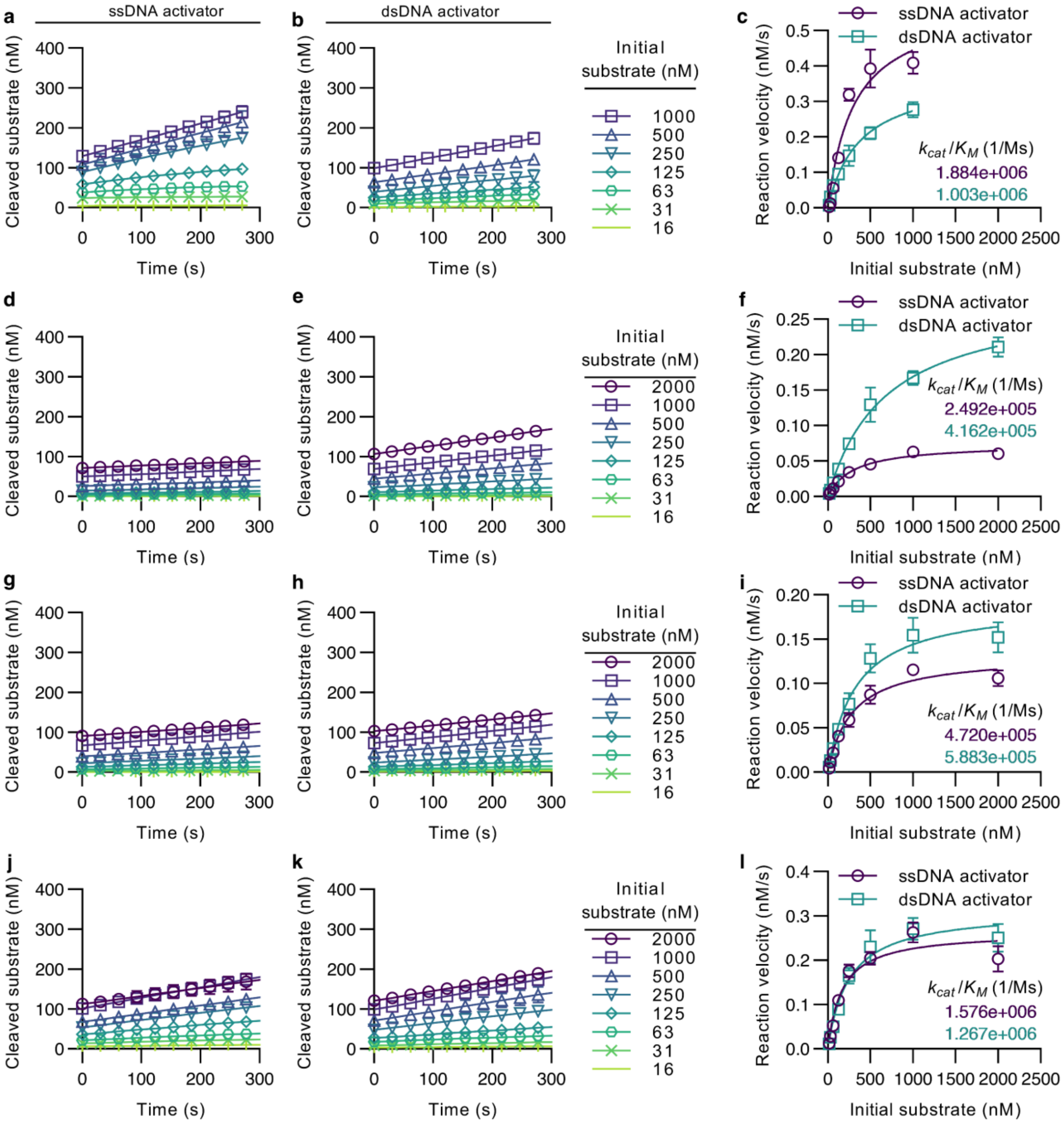

**Fig. S2. LbCas12a trans-cleavage progress curves and Michaelis-Menten kinetics. a** Trans-cleavage progress curves for LbCas12a complexed with gRNA-1 (termed LbCas12a-1) and activated with a ssDNA. **b** Trans-cleavage progress curves for LbCas12a-1 activated with dsDNA. **c** LbCas12a-1 Michaelis-Menten curves. **d-e** Trans-cleavage progress curves for LbCas12a-2 activated respectively with ss- and dsDNA. **f** LbCas12a-2 Michaelis-Menten curves. **g-h** Trans-cleavage progress curves for LbCas12a-3 activated respectively with ss- and dsDNA. **i** LbCas12a-3 Michaelis-Menten curves. **j-k** Trans-cleavage progress curves for LbCas12a-4 activated respectively with ss- and dsDNA. **l** LbCas12a-4 Michaelis-Menten curves. All data here is for a 1 nM activated enzyme concentration.

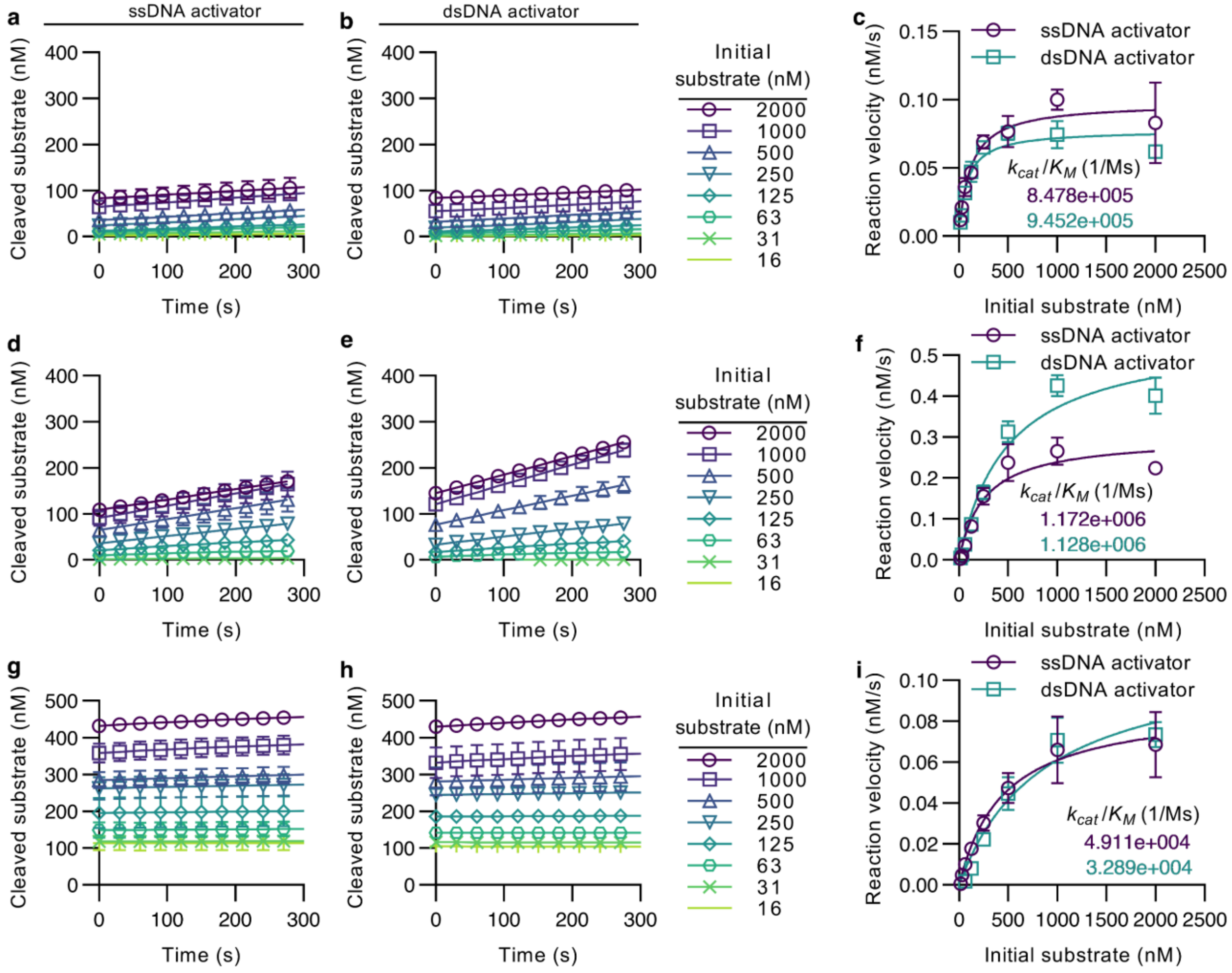

**Fig. S3. Additional LbCas12a trans-cleavage progress curves and Michaelis-Menten kinetics.** **a-b** Trans-cleavage progress curves for LbCas12a-5 activated respectively with ss- and dsDNA. **c** LbCas12a-5 Michaelis-Menten curves. **d-e** Trans-cleavage progress curves for LbCas12a-6 activated respectively with ss- and dsDNA. **f** LbCas12a-6 Michaelis-Menten curves. **g-h** Trans-cleavage progress curves for LbCas12a-7 activated respectively with ss- and dsDNA. **f** LbCas12a-7 Michaelis-Menten curves. Data in **a-f** is for a 1 nM activated enzyme concentration and data in **g-i** is for a 4 nM activated enzyme concentration.

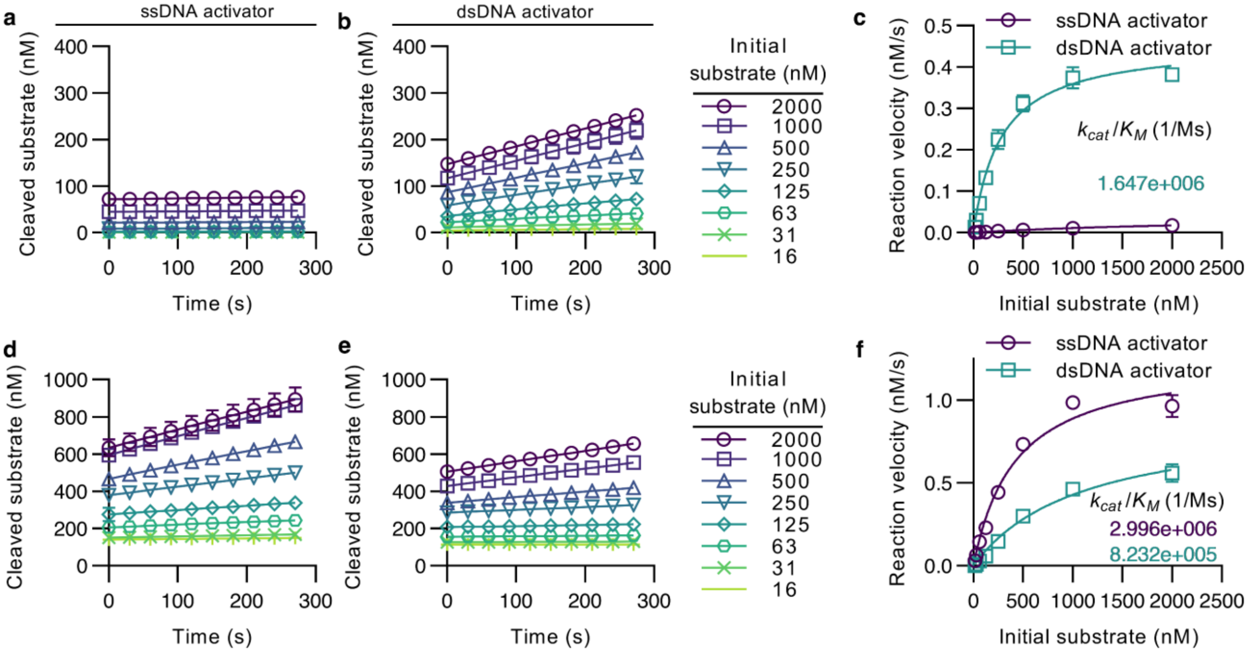

**Fig. S4. AsCas12a trans-cleavage progress curves and Michaelis-Menten kinetics. a-b** Trans-cleavage progress curves for AsCas12a-1 activated respectively with ss- and dsDNA. **c** AsCas12a-1 Michaelis-Menten curves. Note the ssDNA activator for AsCas12a-1 resulted in almost no cleavage. Therefore, no kinetics are reported for this gRNA-activator pair. **d-e** Trans-cleavage progress curves for AsCas12a-2 activated respectively with ss- and dsDNA. **f** AsCas12a-2 Michaelis-Menten curves. All data here is for a 1 nM activated enzyme concentration.

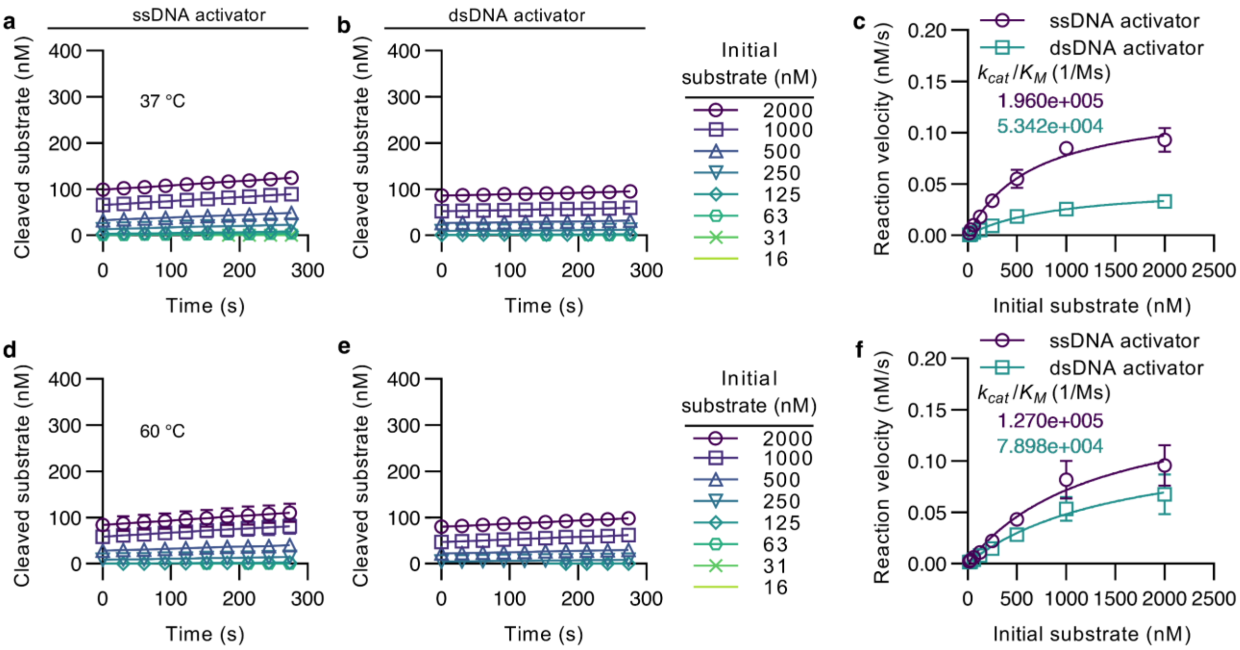

**Fig. S5. AapCas12b trans-cleavage progress curves and Michaelis-Menten kinetics. a-b** Trans-cleavage progress curves for AapCas12b-1 at 37 °C activated respectively with ss- and dsDNA. **c** AapCas12b-1 Michaelis-Menten curves at 37 °C. **d-e** Trans-cleavage progress curves for AapCas12b-1 at 60 °C activated respectively with ss- and dsDNA. **f** AapCas12b-1 Michaelis-Menten curves at 60 °C. All data here is for a 1 nM activated enzyme concentration.

### S3. LbCas12a, AsCas12a, and AapCas12b limits of detection

This section presents measured fluorescence versus time curves for the LbCas12a, AsCas12a, and AapCas12b complexes (**Figs. S6-S9**). We here also show the endpoint fluorescence-based limit of detection (LoD) as well as the maximum trans-cleavage velocity-based LoD.

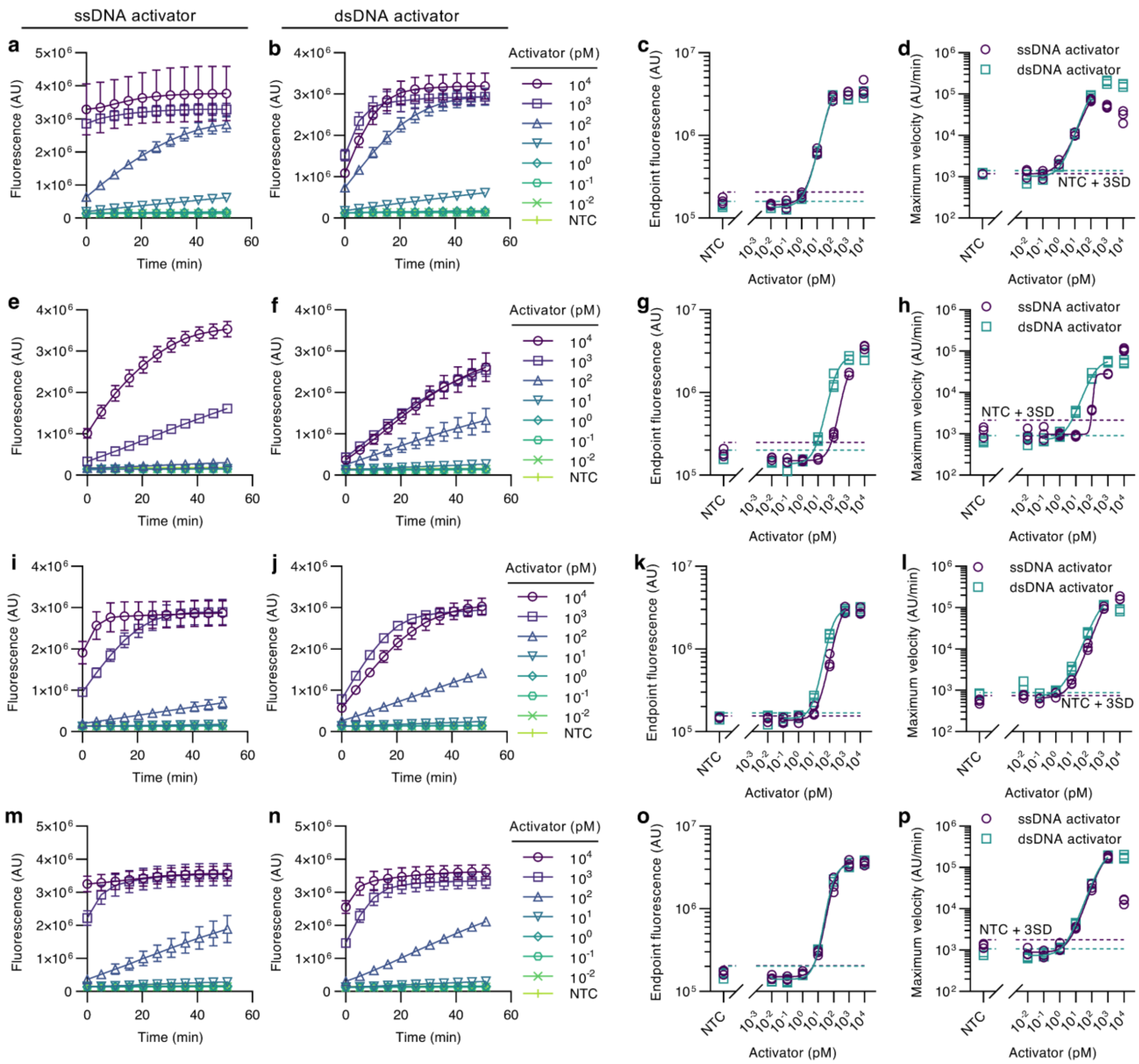

**Fig. S6. LbCas12a limit of detection. a** Fluorescence versus time for LbCas12a-1 activated with a ssDNA. The activator concentration was varied between 0 (NTC) and $10^4$ pM. **b** Fluorescence versus time for LbCas12a-1 activated with dsDNA. **c** Endpoint fluorescence after 1 h for LbCas12a-1. The dashed lines correspond to the detection threshold and their evaluation is shown in the main work (see **Methods**). **d** Maximum trans-cleavage cleavage velocity for LbCas12a-1. Note the maximum velocity decreases at very high activator concentrations because nearly all reporters have been cleaved prior to measurement. **e-f** Fluorescence versus time for LbCas12a-2 activated respectively with ss- and dsDNA. **g** Endpoint fluorescence after 1 h for LbCas12a-2. **h**

Maximum trans-cleavage cleavage velocity for LbCas12a-2. **i-j** Fluorescence versus time for LbCas12a-3 activated respectively with ss- and dsDNA. **k** Endpoint fluorescence after 1 h for LbCas12a-3. **l** Maximum trans-cleavage cleavage velocity for LbCas12a-3. **m-n** Fluorescence versus time for LbCas12a-4 activated respectively with ss- and dsDNA. **o** Endpoint fluorescence after 1 h for LbCas12a-4. **p** Maximum trans-cleavage cleavage velocity for LbCas12a-4. All data here is for a 200 nM DNA reporter concentration.

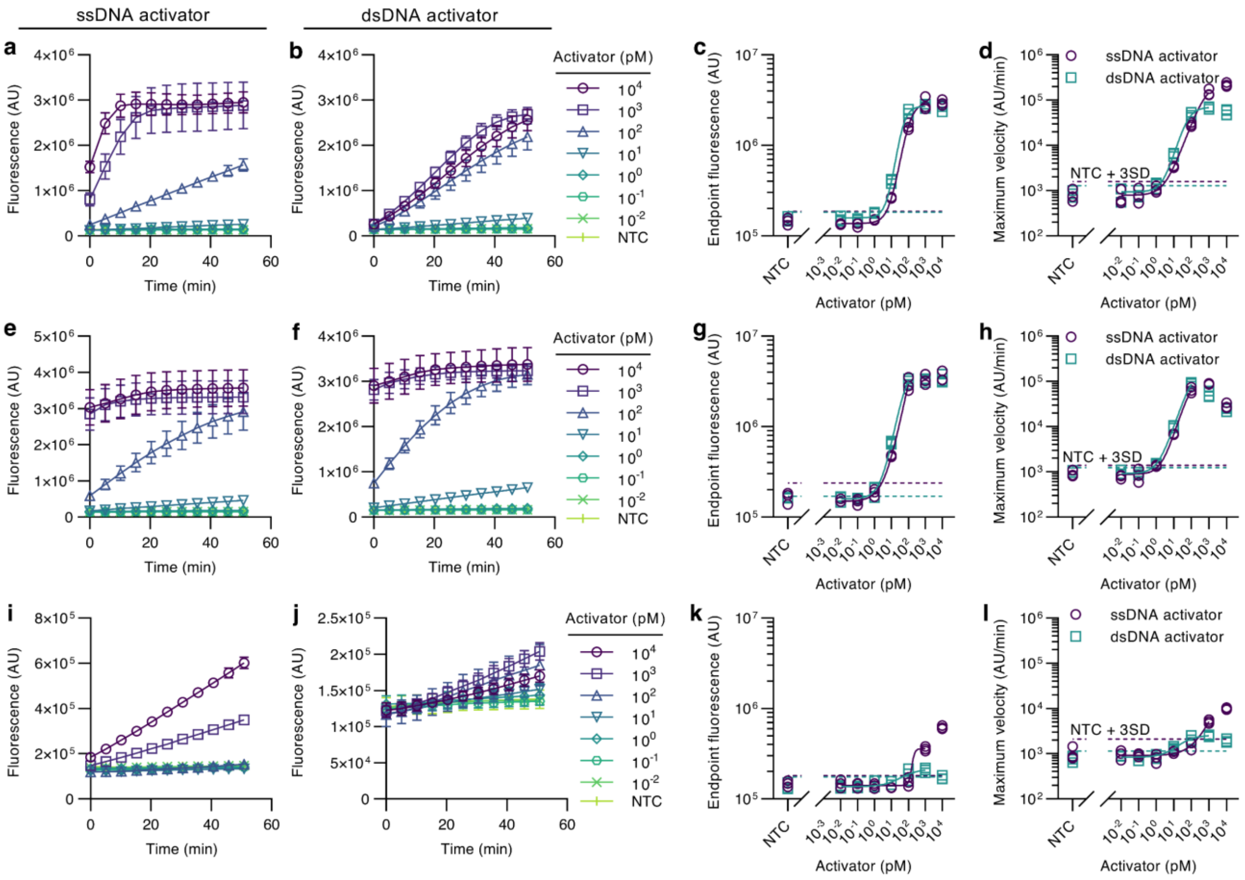

**Fig. S7. Additional LbCas12a limit of detection. a-b** Fluorescence versus time for LbCas12a-5 activated respectively with ss- and dsDNA. **c** Endpoint fluorescence after 1 h for LbCas12a-5. **d** Maximum trans-cleavage cleavage velocity for LbCas12a-5. **e-f** Fluorescence versus time for LbCas12a-6 activated with, respectively, ssDNA and dsDNA. **g** Endpoint fluorescence after 1 h for LbCas12a-6. **h** Maximum trans-cleavage cleavage velocity for LbCas12a-6. **i-j** Fluorescence versus time for LbCas12a-7 activated respectively with ss- and dsDNA. **k** Endpoint fluorescence after 1 h for LbCas12a-7. **l** Maximum trans-cleavage cleavage velocity for LbCas12a-7. All data here is for a 200 nM DNA reporter concentration.

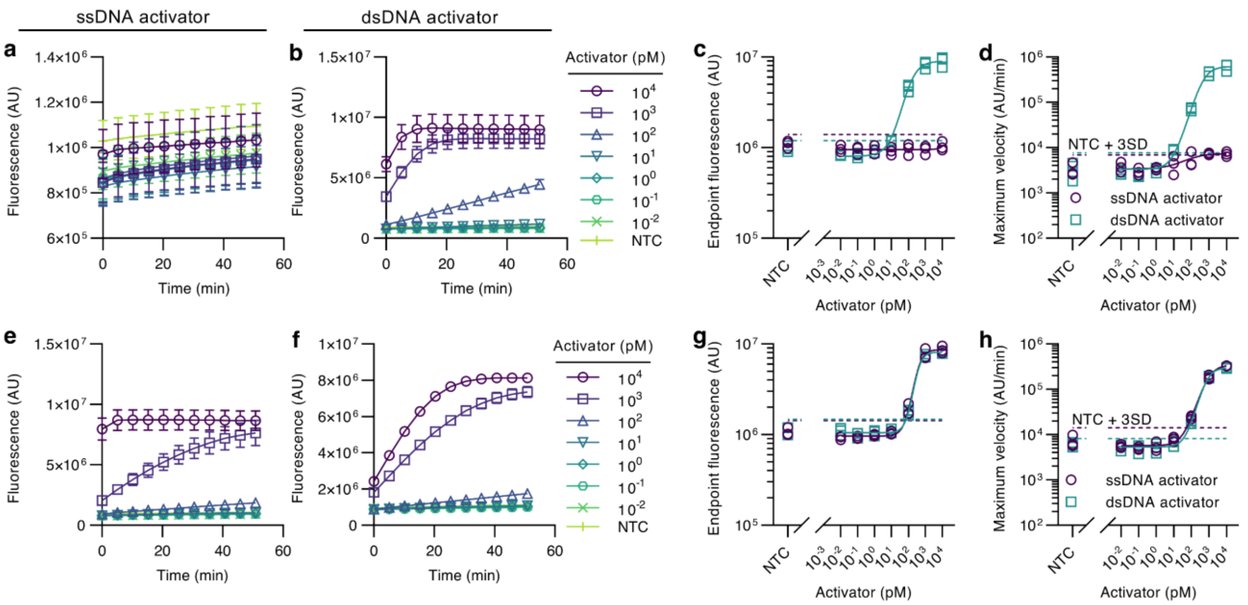

**Fig. S8. AsCas12a limit of detection. a-b** Fluorescence versus time for AsCas12a-1 activated respectively with ss- and dsDNA. **c** Endpoint fluorescence after 1 h for AsCas12a-1. **d** Maximum trans-cleavage cleavage velocity for AsCas12a-1. Note that, as expected from the observed poor cleavage of ssDNA-activated AsCas12a-1 (**Supplementary Fig. S4a**), this enzyme-gRNA-activator combination cannot detect target even at high concentrations (= $10^4$ pM). **e-f** Fluorescence versus time for AsCas12a-2 activated respectively with ss- and dsDNA. **g** Endpoint fluorescence after 1 h for AsCas12a-2. **h** Maximum trans-cleavage cleavage velocity for AsCas12a-2. All data here is for a 200 nM DNA reporter concentration.

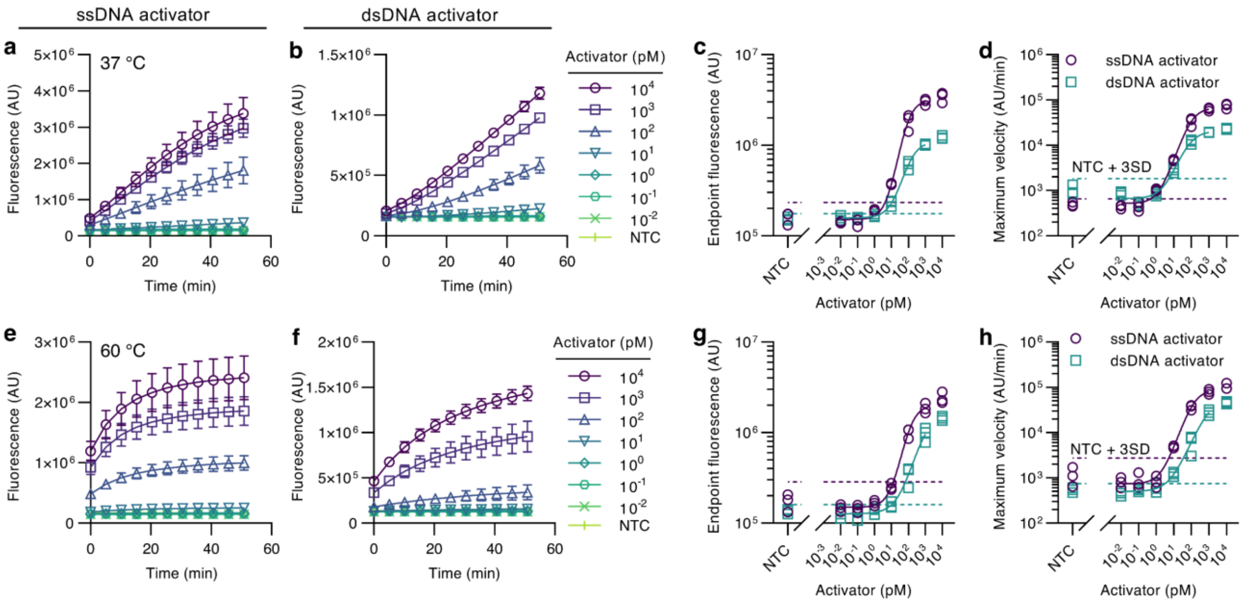

**Fig. S9. AapCas12b limit of detection. a-b** Fluorescence versus time for AapCas12b-1 at 37 °C activated respectively with ss- and dsDNA. **c** Endpoint fluorescence after 1 h at 37 °C for AapCas12b-1. **d** Maximum trans-cleavage cleavage velocity at 37 °C for AsCas12a-1. **e-f** Fluorescence versus time for AapCas12b-1 at 60 °C activated respectively with ss- and dsDNA. **g** Maximum trans-cleavage cleavage velocity at 60 °C for AapCas12b-1. **h** Maximum cleavage velocity in 1 h at 60 °C for AsCas12a-1. All data here is for a 200 nM DNA reporter concentration.

### S4. RNA reporter calibration curves

**Figure S10** shows the calibration curves for the cleaved and uncleaved ssRNA reporters used in this work. Again, these curves were used to quantify cleavage activity in molar units. However, the relatively high reporter concentration necessitated a nonlinear calibration as follows.

First, linear curves of best fit are calculated for reporter concentrations at or below 1 μM. At concentrations around 2 μM, the fluorophore inner filter effect (IFE) is significant, and fluorescence decreases with increasing concentration.[2] We here modeled the effect of the IFE as

$$F = \frac{F_{ucl}c_{ucl} + F_{cl}c_{cl}}{IFE}, \tag{1}$$

where

$$IFE = 10^{\frac{c_{ucl}+c_{cl}}{C_0}} \tag{2}$$

and

$$C_0 = \frac{C_{ucl,0} + C_{cl,0}}{2}. \tag{3}$$

In the latter equations, $F$ is the measured fluorescence, $c_{ucl}$ and $c_{cl}$ are respectively the uncleaved and cleaved reporter concentration, $F_{ucl}$ and $F_{cl}$ are respectively slopes of the best fit lines to the uncleaved and cleaved fluorescence data at $c_{ucl}$, $c_{cl} \leq 1$ μM (where the IFE is negligible), and $C_{ucl,0}$ and $C_{cl,0}$ were respectively determined from the power law curves of best fit to the uncleaved and cleaved fluorescence data. Eq. (1) can be rearranged to calculate $c_{cl}$ as

$$c_{cl} = \frac{F10^{\frac{c_{ucl}+c_{cl}}{C_0}} - F_{ucl}c_0}{F_{cl} - F_{ucl}}, \tag{4}$$

where $c_0$ is the initial uncleaved reporter concentration. Hence, the IFE-inclusive calibration curves were used to quantify cleavage activity in molar units for Cas13-based RNA detection.

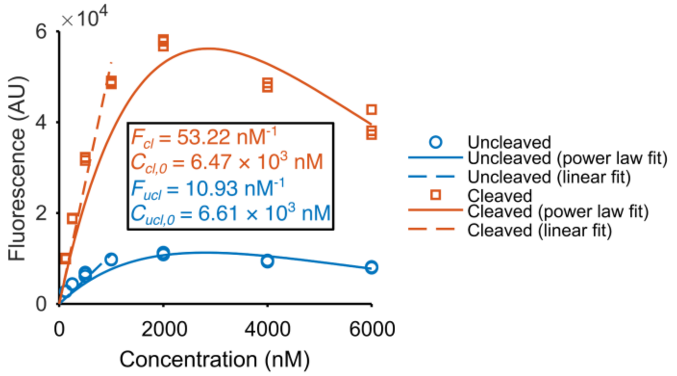

**Fig. S10. RNA reporter calibration curves.** Measured fluorescence versus concentration of uncleaved and cleaved FAM-BHQ RNA reporters. Lines (dashed line) and power law curves (solid lines) of best fit to the experimental data were obtained by linear regression. Data was taken in an Infinite® 200 PRO (Tecan Group Ltd., Switzerland).

### S5. LwaCas13a and LbuCas13a Michaelis-Menten kinetics

This section presents the linear portion of the trans-cleavage progress curves for the LwaCas13a and LbuCas13a complexes (**Figs. S11** and **S12**). Also presented are the Michaelis-Menten fits to the initial reaction velocity to extract relevant kinetic parameters.

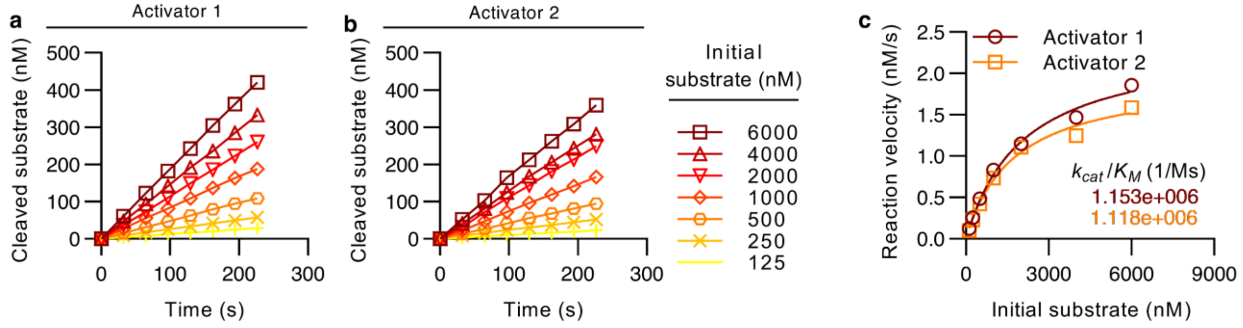

**Fig. S11. LwaCas13a trans-cleavage progress curves and Michaelis-Menten kinetics. a** Trans-cleavage progress curves for LwaCas13a-1 activated with a complementary RNA. **b** Trans-cleavage progress curves for LwaCas13a-2 activated with a complementary RNA. **c** Michaelis-Menten curves for LwaCas13a-1 and LwaCas13a-2.

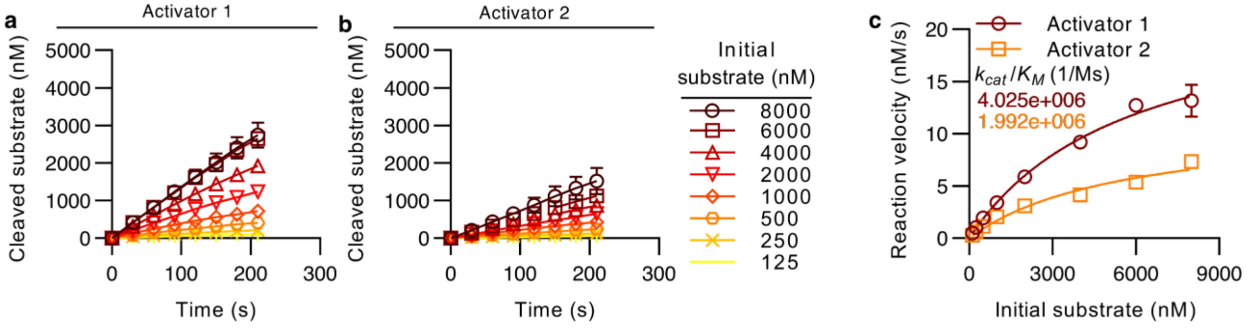

**Fig. S12. LbuCas13a trans-cleavage progress curves and Michaelis-Menten kinetics. a** Trans-cleavage progress curves for LbuCas13a-1 activated with a complementary RNA. **b** Trans-cleavage progress curves for LbuCas13a-2 activated with a complementary RNA. **c** Michaelis-Menten curves for LbuCas13a-1 and LbuCas13a-2.

### S6. LwaCas13a and LbuCas13a limits of detection

This section presents fluorescence measurements versus time curves for the LwaCas13a and LbuCas13a complexes (**Figs. S13** and **S14**). We here also show the endpoint fluorescence-based LoD as well as the maximum trans-cleavage velocity-based LoD.

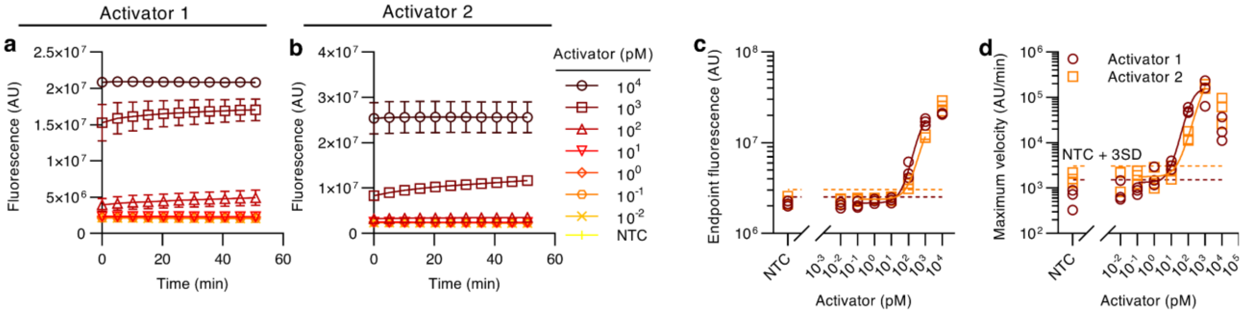

**Fig. S13. LwaCas13a limit of detection. a** Fluorescence versus time for LwaCas13a-1 activated with a complementary RNA. **b** Fluorescence versus time for LwaCas13a-2 activated with a complementary RNA. **c** Endpoint fluorescence after 1 h for LwaCas13a-1 and LwaCas13a-2. **d** Maximum trans-cleavage velocity for LwaCas13a-1 and LwaCas13a-2. All data here is for a 1 µM RNA reporter concentration.

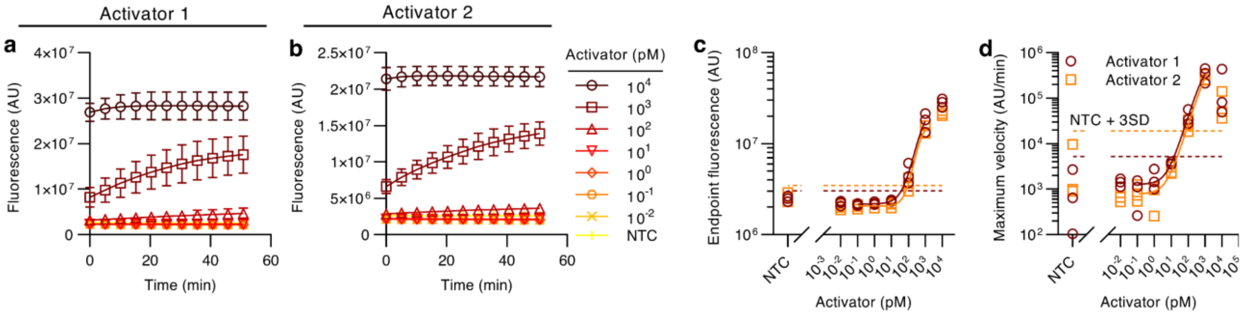

**Fig. S14. LbuCas13a limit of detection. a** Fluorescence versus time for LbuCas13a-1 activated with a complementary RNA. **b** Fluorescence versus time for LbuCas13a-2 activated with a complementary RNA. **c** Endpoint fluorescence after 1 h for LbuCas13a-1 and LbuCas13a-2. **d** Maximum trans-cleavage velocity for LbuCas13a-1 and LbuCas13a-2. All data here is for a 1 µM RNA reporter concentration.

### S7. Cas13a correlations

This section presents plots showing the relation between $L_E$ and $P_E$ as well as $L_V$ and $P_V$ for LwaCas13a and LbuCas13a (**Fig. S15**). We also present here our method to quantify overall agreement between measured kinetic parameters and LoDs using the heuristic quantity $H$ applied to LwaCas13a and LbuCas13a.

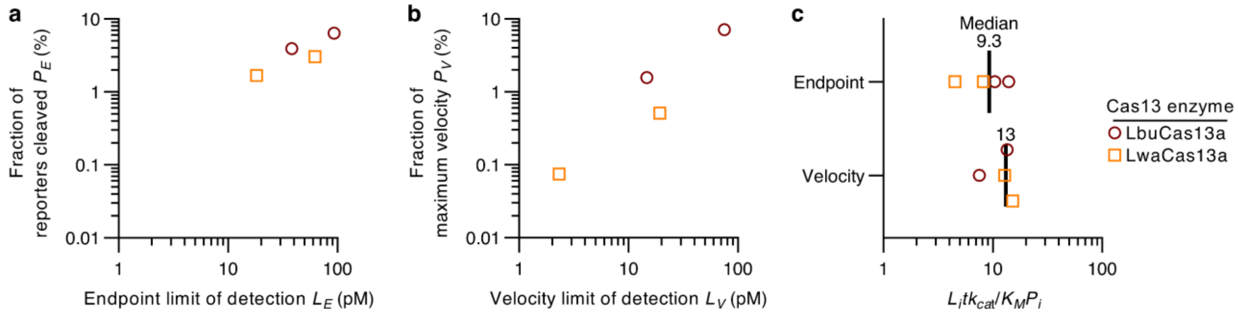

**Fig. S15. Cas13a correlations. a** Fraction of cleaved reporters at the endpoint LoD ($P_E$) versus endpoint-based LoD ($L_E$) for LbuCas13a and LwaCas13a. **b** Fraction of maximum cleavage velocity across all target concentrations ($P_V$) versus velocity-based LoD ($L_V$) for the same Cas13-gRNA-target combinations. **c** Heuristic $H = L_i t k_{cat}/K_M P_i$ evaluated for $i = E$ (endpoint) and $V$ (velocity).

### S8. Michaelis-Menten kinetic parameters with confidence intervals

**Table S1** summarizes confidence intervals for the measured kinetic parameters. Confidence intervals were estimated in GraphPad Prism 9 (GraphPad Software, CA, USA) using the asymmetrical (profile-likelihood option).

**Table S1. Summary of experimentally measured kinetic parameters.** This table complements the main work (**Table 1**), providing 95% confidence intervals on $k_{cat}$ and $K_M$ measurements.

| | | 95% confidence intervals | | |
|---|---|---|---|---|
| Name | Target | $k_{cat}$ ($s^{-1}$) | $K_M$ (nM) | Ref. |
| LbCas12a-1 | ssDNA | 0.49 to 0.71 | 200 to 490 | 3 |
| | dsDNA | 0.33 to 0.43 | 290 to 500 | |
| LbCas12a-2 | ssDNA | 0.068 to 0.081 | 230 to 380 | |
| | dsDNA | 0.26 to 0.32 | 550 to 860 | |
| LbCas12a-3 | ssDNA | 0.12 to 0.14 | 220 to 370 | |
| | dsDNA | 0.17 to 0.21 | 240 to 440 | |
| LbCas12a-4 | ssDNA | 0.24 to 0.30 | 130 to 270 | 4 |
| | dsDNA | 0.28 to 0.35 | 190 to 360 | |
| LbCas12a-5 | ssDNA | 0.086 to 0.11 | 73 to 180 | 5 |
| | dsDNA | 0.071 to 0.085 | 58 to 110 | |
| LbCas12a-6 | ssDNA | 0.26 to 0.35 | 170 to 390 | 1 |
| | dsDNA | 0.49 to 0.64 | 350 to 700 | |
| LbCas12a-7 | ssDNA | 0.019 to 0.027 | 290 to 720 | 6 |
| | dsDNA | 0.023 to 0.036 | 560 to 1400 | |
| AsCas12a-1 | ssDNA | - | - | 7 |
| | dsDNA | 0.43 to 0.49 | 230 to 340 | |
| AsCas12a-2 | ssDNA | 1.2 to 1.4 | 330 to 540 | 8 |
| | dsDNA | 0.78 to 1.0 | 830 to 1500 | |
| AapCas12b-1 | ssDNA | 0.12 to 0.15 | 500 to 880 | 9 |
| | dsDNA | 0.045 to 0.054 | 750 to 1100 | |
| | ssDNA* | 0.13 to 0.23 | 770 to 2400 | |
| | dsDNA* | 0.090 to 0.21 | 840 to 3600 | |
| LwaCas13a-2 | RNA | 2.2 to 2.6 | 1700 to 2500 | 7 |
| LwaCas13a-2 | RNA | 1.8 to 2.1 | 1500 to 2200 | |
| LbuCas13a-1 | RNA | 20 to 28 | 4400 to 7900 | 10 |

| | | | |
|---|---|---|---|
| LbuCas13a-2 | RNA | 8.5 to 17 | 3200 to 12000 |

* Trans-cleavage experiments performed at 60 °C.

### S9. Lists of oligos used in this work

This section presents all oligos used in this work, including all gRNAs (**Table S2**), DNA and RNA activators (**Table S3**), and DNA and RNA reporters (**Table S4**).

**Table S2. List of gRNAs used in this work.** All oligos here were purchased from GeneLink, Inc. (USA) except for the gRNA for LbCas12a-4 which was purchased from Integrated DNA Technologies (IDT), Inc. (USA).

| gRNA | Sequence (5'-3') | Ref. |
|---|---|---|
| LbCas12a-1 | UAA UUU CUA CUA AGU GUA GAU GUG GUA UUC UUG CUA GUU AC | 3 |
| LbCas12a-2 | UAA UUU CUA CUA AGU GUA GAU CCC CCA GCG CUU CAG CGU UC | |
| LbCas12a-3 | UAA UUU CUA CUA AGU GUA GAU AAU UAC UUG GGU GUG ACC CU | |
| LbCas12a-4 | UAA UUU CUA CUA AGU GUA GAU CUC AGG GCG GAC UGG GUG CU | 4 |
| LbCas12a-5 | UAA UUU CUA CUA AGU GUA GAU CCG CGG GUG GUC CCG GAC AG | 5 |
| LbCas12a-6 | UAA UUU CUA CUA AGU GUA GAU UGU AUG GCA UGA GUA ACG AA | 1 |
| LbCas12a-7 | UAA UUU CUA CUA AGU GUA GAU CGU CGC CGU CCA GCU CGA CC | 6 |
| AsCas12a-1 | UAA UUU CUA CUC UUG UAG AUC UGU GUU UAU CCG CUC ACA A | 7 |
| AsCas12a-2 | GGU UUA AUU UCU ACU CUU GUA GAU CCG GCA AGC UGC CCG UGC CC | 8 |
| AapCas12b-1 | GUC UAG AGG ACA GAA UUU UUC AAC GGG UGU GCC AAU GGC CAC UUU CCA GGU GGC AAA GCC CGU UGA GCU UCU CAA AUC UGA GAA GUG GCA CCG AAG AAC GCU GAA GCG CUG | 9 |
| LwaCas13a-1 | GAT TTA GAC TAC CCC AAA AAC GAA GGG GAC TAA AAC TGC TTC TGT CCA GTG AGC ATG GTC TTC G | 7 |
| LwaCas13a-2 | GAT TTA GAC TAC CCC AAA AAC GAA GGG GAC TAA AAC ACT CCC TAG AAC CAC GAC AGT TTG CCT T | |
| LbuCas13a-1 | GAC CAC CCC AAA AAU GAA GGG GAC UAA AAC GGU CCA CCA AAC GUA AUG CG | 10 |
| LbuCas13a-2 | GAC CAC CCC AAA AAU GAA GGG GAC UAA AAC UUU GCG GCC AAU GUU UGU AA | |

**Table S3. List of activators used in this work.** List includes target and non-target strands (TS and NTS, respectively). All Cas12 oligos are DNA while all Cas13 oligos are RNA. Cas12 oligos were purchased from Elim Biopharmaceuticals, Inc. (USA). Cas13 oligos were purchased from GeneLink, Inc. (USA). dsDNA activators were prepared by annealing TS and NTS (see **Methods** in the main work).

| Activator | | Sequence (5'-3') | Ref. |
|---|---|---|---|
| LbCas12a-1 | TS | ATC GAA GCG CAG TAA GGA TGG CTA GTG TAA CTA GCA AGA ATA CCA CGA AAG CAA GAA AAA | 3 |
| | NTS | TTT TTC TTG CTT TCG TGG TAT TCT TGC TAG TTA CAC TAG CCA TCC TTA CTG CGC TTC GAT | |
| LbCas12a-2 | TS | ACA TTC CGA AGA ACG CTG AAG CGC TGG GGG CAA ATT GTG C | |
| | NTS | GCA CAA TTT GCC CCC AGC GCT TCA GCG TTC TTC GGA ATG T | |
| LbCas12a-3 | TS | CCG AGT CTT CAG GGT CAC ACC CAA GTA ATT GAA AAG ACA C | |
| | NTS | GTG TCT TTT CAA TTA CTT GGG TGT GAC CCT GAA GAC TCG G | |
| LbCas12a-4 | TS | CCA CTA CCT GAG CAC CCA GTC CGC CCT GAG CAA AGA CCC C | 4 |
| | NTS | GGG GTC TTT GCT CAG GGC GGA CTG GGT GCT CAG GTA GTG G | |
| LbCas12a-5 | TS | ATC AGC GAT CGT GGT CCT GCG GGC TTT GCC GCG GGT GGT CCC GGA CAG GCC GAG TTT GGT CAT CAG CCG TTC | 5 |
| | NTS | GAA CGG CTG ATG ACC AAA CTC GGC CTG TCC GGG ACC ACC CGC GGC AAA GCC CGC AGG ACC ACG ATC GCT GAT | |
| LbCas12a-6 | TS | ATT ATA TTC TTC GTT ACT CAT GCC ATA CAT AAA TGG ATA GA | 1 |
| | NTS | TCT ATC CAT TTA TGT ATG GCA TGA GTA ACG AAG AAT ATA AT | |
| LbCas12a-7 | TS | GCC GGG GTG GTG CCC ATC CTG GTC GAG CTG GAC GGC GAC GTA AAC GGC CAC AAG C | 6 |
| | NTS | GCT TGT GGC CGT TTA CGT CGC CGT CCA GCT CGA CCA GGA TGG GCA CCA CCC CGG C | |
| AsCas12a-1 | TS | GGC CAG TGA ATT CGA GCT CGG TAC CCG GGG ATC CTC TAG AAA TAT GGA TTA CTT GGT AGA ACA GCA ATC TAC TCG ACC TGC AGG CAT GCA AGC TTG GCG TAA TCA TGG TCA TAG CTG TTT CCT GTG TTT ATC CGC TCA CAA TTC CAC ACA ACA TAC GAG CCG GAA GCA TAA AG | 7 |
| | NTS | CTT TAT GCT TCC GGC TCG TAT GTT GTG TGG AAT TGT GAG CGG ATA AAC ACA GGA AAC AGC TAT GAC CAT GAT TAC GCC AAG CTT GCA TGC CTG CAG GTC | |

| | | | |
|---|---|---|---|
| | | GAG TAG ATT GCT GTT CTA CCA AGT AAT CCA TAT TTC TAG AGG ATC CCC GGG TAC CGA GCT CGA ATT CAC TGG CC | |
| AsCas12a-2 | TS | TAG GGC ACG GGC AGC TTG CCG GTA AAG TCA CTA TGC C | 8 |
| | NTS | GGC ATA GTG ACT TTA CCG GCA AGC TGC CCG TGC CCT A | |
| AapCas12b-1 | TS | AAA TTG CAC AAT TTG CCC CCA GCG CTT CAG CGT TCT TCG GAA TGT CGC GCA TTG GCA TGG | 9 |
| | NTS | CCA TGC CAA TGC GCG ACA TTC CGA AGA ACG CTG AAG CGC TGG GGG CAA ATT GTG CAA TTT | |
| LwaCas13a-1 | TS | GGC CAA CCU CUC AAC AAU GAC GAA GAC CAU GCU CAC UGG ACA GAA GCA AAA AUG CUG CUG GAC AAC AU | 7 |
| LwaCas13a-2 | TS | UUC AAG GAC GCA CAU GCC AAA AGG CAA ACU GUC GUG GUU CUA GGG AGU CAA GAA GGA GCA GUU CAC AC | |
| LbuCas13a-1 | TS | AAA UGC ACC CCG CAU UAC GUU UGG UGG ACC CUC AGA UUC A | 10 |
| LbuCas13a-2 | TS | CUA AUC AGA CAA GGA ACU GAU UAC AAA CAU UGG CCG CAA AUU GCA CAA UUU GCC CCC AGC | |

**Table S4. List of reporters used in this work.** ssDNA reporters were used for all Cas12 experiments and were purchased from the Stanford Protein and Nucleic Acid Facility (PAN). ssRNA reporters were used for all Cas13 experiments and were purchased from GeneLink, Inc. (USA).

| Reporter | Sequence (5’-3’) |
|---|---|
| ssDNA | [5-Fam] TTA TTA TT [3-BHQ-1] |
| ssRNA | [5-Fam] UUU UU [3-BHQ-1] |
